# Distinct iron metabolism regulation during cachexia progression in murine models of intestinal and pancreatic cancer

**DOI:** 10.64898/2026.10.02.756162

**Authors:** Auffret Maëlys, Gasrel Pauline, Orfila Luz, Martin Brice, Martin Agnès, Freyssenet Damien, Island Marie-Laure, Ropert Martine, Rébillard Amélie, Derbré Frédéric

## Abstract

**Background:** Cancer cachexia is a multifactorial syndrome characterized by substantial metabolic disturbances including systemic inflammation, and muscle atrophy, which may affect the regulation of iron metabolism in patients. This study aimed to characterize this regulation in two murine models of colorectal and pancreatic cancer, both associated with a high prevalence of cachexia.

**Methods:** Histological, biochemical and molecular analyses were performed in serum, skeletal muscle, liver, spleen and tumor from UN-KC-6141 pancreatic tumor bearing mice and from *Apc*^Min/+^ mice (colorectal cancer model), both models exhibiting variable cachexia severity.

**Results:** Both models induce cachexia, but diverge markedly in their regulation of iron metabolism. UNKC mice display reduced plasma iron availability (-9.6% transferrin saturation, p<0.001) without concomitant anemia, while hepatic hepcidin synthesis is upregulated (+100.6%, p<0.001) by inflammation, promoting ferroportin (FPN) downregulation (-42.4%, p<0.001) and iron sequestration in the spleen (+134% total iron content, p<0.001). In the atrophied gastrocnemius, iron concentration is preserved through increased ferritin (+86.6%, p=0.010) and hemoproteins (i.e., myoglobin, CYSC, COXIV) expression. Conversely, *Apc*^Min/+^ mice exhibit anemia together with hepatic and splenic iron depletion (-48.4%, p<0.001 and - 84.9%, p<0.001, respectively). Hepcidin synthesis is downregulated (-99.9%, p<0.001) despite persistent inflammation, indicating that hypoxia and/or iron-deficiency signals probably predominate over the inflammatory pathway. FPN is markedly decreased (-94.2%, p<0.001) in the iron-depleted spleen despite low hepcidin. In the atrophied quadriceps of *Apc*^Min/+^ mice, iron deficiency is associated with decreased FPN and ferritin (-87.9%, p<0.001, and -38.4%, p=0.016, respectively), and increased TfR1 expression (+305%, p=0.107). Finally, tumor iron import may play a significant role in cachexia progression, as suggested by the association between tumor weight, tumor TfR1 protein levels and body weight loss in UNKC mice.

**Conclusion:** These findings reveal cancer type-specific signatures of iron regulation in cachexia, which appear to be more driven by cancer type specific features than by cachexia severity itself, with skeletal muscle iron metabolism impaired mainly when cachexia is associated with anemia. This suggests that the clinical management of iron metabolism in cachectic patients should be guided by the underlying cancer type.

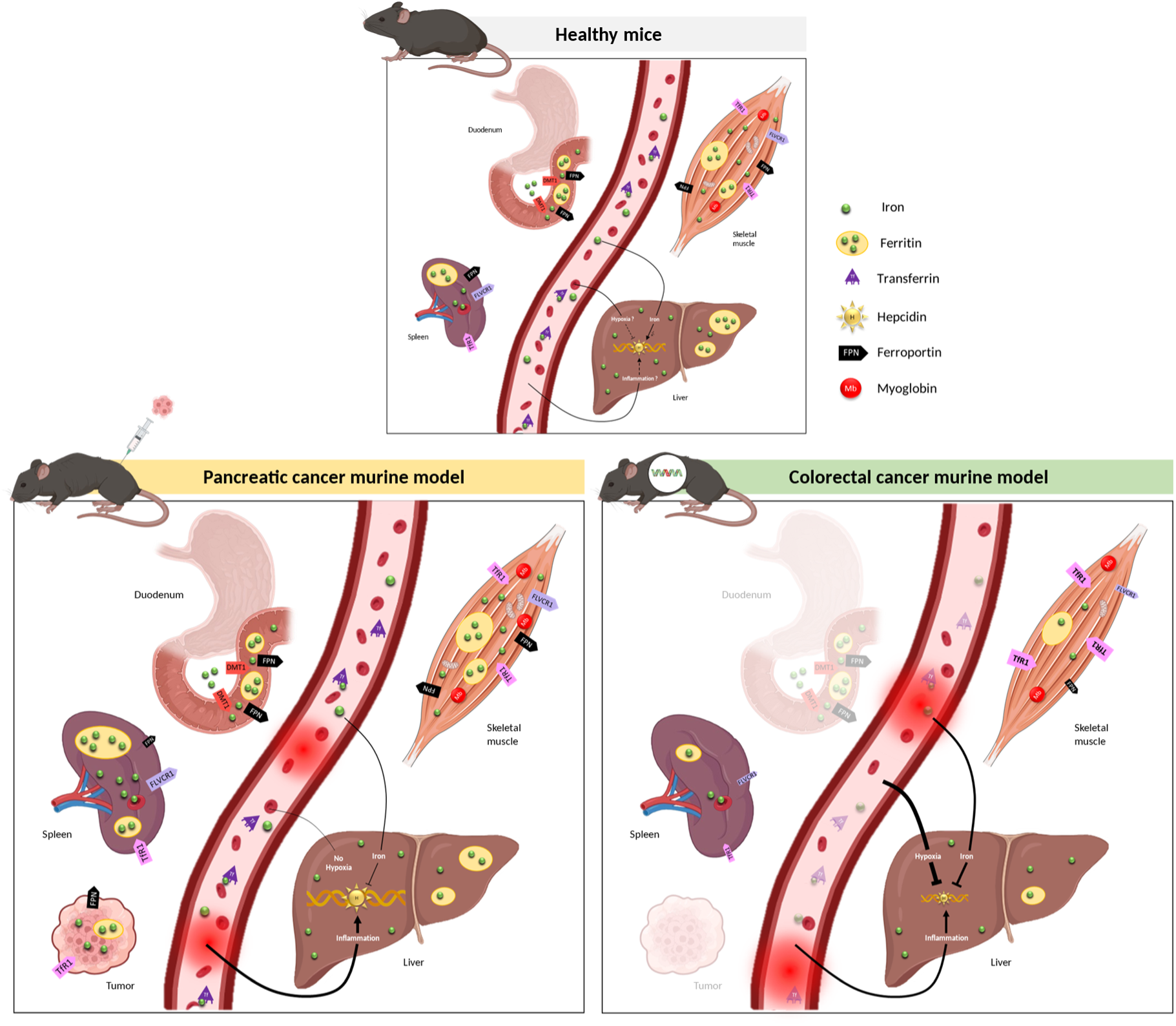

**Abstract figure legend:** Cancer cachexia is associated with iron redistribution, diverging between UN-KC-6141 pancreatic cancer model and the *Apc*^Min/+^ colorectal cancer model. In pancreatic cancer, hepatic hepcidin is upregulated, likely in response to inflammation, and induces splenic iron sequestration, resulting in decreased plasma iron availability. In colorectal cancer, cachexia is associated with anemia together with splenic and hepatic iron depletion, whose signals probably drive hepcidin downregulation despite inflammation. Whatever the model, skeletal muscle appears to engage cellular adaptive regulatory mechanisms to preserve itself from the systemic iron dysregulation induced by cancer cachexia and its associated features.

## INTRODUCTION

Cancer cachexia is a complex multifactorial syndrome characterized by body weight loss and muscle wasting, with or without loss of fat mass, associated with metabolic disturbances, particularly systemic inflammation^1–3^. It affects 50 to 80% of cancer patients, contributing to anorexia, physical deconditioning and fatigue, and ultimately impairing quality of life, anticancer treatment response and survival. Its prevalence is particularly high in digestive cancers, pancreatic cancer being the most frequently associated with this comorbidity, followed by gastrointestinal cancers (i.e., esophageal, gastric and colorectal cancers)^4^. The development of this syndrome results from chronic systemic inflammation triggered by interactions between tumor-derived factors and cytokines released by the host immune system. These persistent inflammatory signals alter numerous central and peripheral signaling pathways, promoting anorexia, hypermetabolism, malabsorption, and loss of fat and muscle mass.^5^ Anemia can also be associated with cancer cachexia in some cases, and be particularly pronounced in gastrointestinal cancers prone to chronic tumor-related bleeding, such as colorectal cancer^6,7^.

Inflammation and muscle atrophy are two hallmark features of cachexia which, independently of this pathological context, are known to modulate iron homeostasis. Yet iron is a micronutrient essential for numerous physiological processes, including energy production through oxidative metabolism, as well as oxygen transport and storage through hemoglobin and myoglobin^8,9^. Iron is therefore particularly abundant in blood and skeletal muscle, and its distribution and metabolism are tightly regulated at cellular and systemic levels in order to prevent both iron deficiency or overload^10^. At the cellular level, this regulation is ensured by the iron-responsive element – iron regulatory protein (IRE-IRP) system, which modulates the translation of iron transport and storage proteins according to cellular needs. Specifically, the IRE-IRP system promotes iron export via the sole iron exporter, ferroportin (FPN), and iron storage via ferritin (FtH) in case of cellular iron excess, while conversely upregulating iron import via TfR1 when the cellular iron pool is deficient^11^. The heme exporter FLVCR1 may also contribute to cellular iron homeostasis, potentially in myoglobin-rich skeletal muscle fibers^12,13^. At systemic level, hepcidin is the key regulator of iron metabolism: once synthesized, it reduces intestinal iron absorption and organ iron release by inducing the degradation and/or inactivation of membrane-bound FPN^14–16^. Hepcidin synthesis is itself regulated by inflammation, hypoxia and iron status, and can be either inhibited or enhanced according to the iron needs of the organism^17,9,18^.

A pro-inflammatory state is therefore an activator of hepcidin synthesis, which is generally associated with decreased plasma iron availability and increased cellular iron sequestration^17,18^. Pathophysiological contexts associated with muscle atrophy, such as age-related sarcopenia and muscle disuse are characterized by iron redistribution between organs, but also within subcellular compartments, consecutive to the loss of muscle mass and to the associated pro-inflammatory state^19–24^. Specifically, elevated iron concentrations in atrophied skeletal muscle are reported in age-related sarcopenia^19–21^, whereas muscle disuse (i.e., bedrest, hindlimb unloading) often induces modulations of plasma iron availability along with an increase of splenic, hepatic and muscle concentrations^22–25^. Finally, in the case of cachectic cancers associated with anemia, red blood cell depletion constitutes an additional factor capable of considerably influence iron homeostasis.

Given the systemic metabolic context of cachexia and the coexistence of several factors each independently able to affect iron metabolism, substantial iron dysregulation is expected during cachexia development. However, systemic iron redistribution under cancer cachexia conditions remains very poorly characterized. The aim of the present study was therefore to characterize the impact of cachectic cancers on systemic iron regulation and distribution, and to describe the organ crosstalk involved. To this end, we explored systemic and cellular iron metabolism in two murine models of cachectic cancer : the UN-KC-6141 model of pancreatic cancer, and the *Apc*^Min/+^ model of colorectal cancer associated with tumor bleeding likely to induce anemia^26^.

## MATERIAL AND METHODS

### Animal models

Animal experiments received approval from the Ethics Committee for Animal Experimentation (authorizations APAFIS#23633 and APAFIS#24956) in accordance with the European Community guidelines (Directive 86/609/EEC). Mice were housed in a temperature-controlled room (21 ± 2°C) with a 12-h:12-h light dark cycle and provided with standard rodent chow and water *ad libitum*.

For UN-KC-6141 tumor model, 30 6-week-old male C57BL/6 (Janvier Labs, Le Genest St Isle, France) were housed in separate cages and were randomly assigned to two groups after a week of acclimatization. Mice received either a subcutaneous injection of 1×10^6^ UN-KC-6141 cancer cells (UNKC, n=20) or PBS (CTL, n=10). Mice were monitored daily for body weight and food intake. After 34 days following the injection, mice were deeply anesthetized with isoflurane (2-3%). Intracardiac blood was collected in dry tubes, and the mice were euthanized by cervical dislocation. Liver, spleen, skeletal muscles (i.e., soleus, gastrocnemius, quadriceps) and tumor were dissected, weighed and then either frozen in liquid nitrogen or fixed in 4% paraformaldehyde (PFA). Microhematocrit tubes were filled from blood samples for hematocrit (Hct) determination by centrifugation (3000 RPM, 10 min, RT). The blood was then centrifuged (1500 g, 10 min, 4°C) for serum sampling. Mice found dead before the end of the protocol (n=1 in UNKC) or mice exhibiting no tumor growth or no cachexia (n = 6 in UNKC) were excluded from the study.

Tissue samples from *Apc*^Min/+^ mice originated from an experiment conducted at the Inter-University Laboratory of Human Movement Biology (Saint-Étienne, France) and reported by Martin et al., 2022^27^. As previously described, *Apc* mice were maintained on a C57BL/6J background and paired with corresponding healthy control littermates. 11- to 23-week-old control and *Apc* mice were anaesthetized (90 mg/kg ketamine and 10 mg/kg xylazine), organs were removed and weighed, and mice were euthanized by cervical dislocation. Tissue samples from 24 *Apc*^Min/+^ mice (*Apc*, n=24) and 24 control mice (CTL, n=24) were selected depending on age and cachexia severity features for the present study. In both models, data analyses were subsequently performed without blinding to group allocation.

### Degree of cancer induced cachexia severity

Cachexia severity was categorized based on final body weight (excluding tumor mass), quadriceps, gastrocnemius and soleus muscles mass, and adipose tissue mass (visceral, epididymal, retroperitoneal and perirenal). Categories were defined as mild, moderate, or severe, based on standard deviations from healthy control mice values. Mice were categorized as having mild cachexia when their variables values were within 1 SD (Standard Deviation) of the control mean, moderate cachexia when values were between 1 SD and 2 SDs, and severe cachexia when values were more than 2SDs below the control mean^28,29^.

### Blood analyses and serum assays

Serum iron and unsaturated iron-binding capacity (UIBC) were measured using a Cobas 8000 analyzer (Roche®) with the Cobas® reagents 03183696 122 and 04536355 190, respectively (Roche Diagnostics, Meylan, France) in the biochemistry laboratory of Rennes Pontchaillou Hospital. Serum transferrin saturation was calculated as [serum iron/ (serum iron + UIBC)] × 100. Plasma IL-6 levels and serum hepcidin concentrations were measured using specific ELISA kits (LEGEND MAX™ Mouse IL-6 ELISA Kit, BioLegend and S-1465 Mouse Hepcidin-25 ELISA Kit BMA Biomedicals, respectively) following the manufacturer’s protocol. IL-6 concentrations were calculated based on the standard curve generated during each assay run, whereas hepcidin concentrations were determined using the manufacturer’s theoretical reference curve.

### Iron quantifications in liver, spleen, skeletal muscles and tumor

Iron concentrations in the liver, spleen, gastrocnemius, quadriceps and tumor were quantified at the AEM2 Platform (Université de Rennes - CHU Rennes) by inductively coupled plasma mass spectrometry on an X-Series IO from ThermoFisher Scientific equipped with collision cell technology (AEM2 platform UF3142), as previously described^30^.

### Western Blot analyses

Tissues were ground into powder in liquid nitrogen and used for protein or RNA extraction. Cytosolic protein extraction was performed and sample protein concentrations were determined as previously described^30^. Proteins were separated by SDS-PAGE (8 to 14% depending on the target protein molecular weight), transferred onto nitrocellulose membranes (Trans-Blot Turbo Transfer System, Bio-Rad), and incubated with the appropriate primary and secondary antibodies (Table 1). All blots were scanned using the Odyssey Imaging System (LI-COR), and band signals were quantified with ImageStudioLite software version 5.2.5 (LI-COR Biosciences). All blots were corrected for loading based on total protein stain signal (Revert™ 700 Total Protein Stain, LI-COR Biosciences) and normalized to the mean signal of the corresponding control group for each experimental model.

**Table 1.**
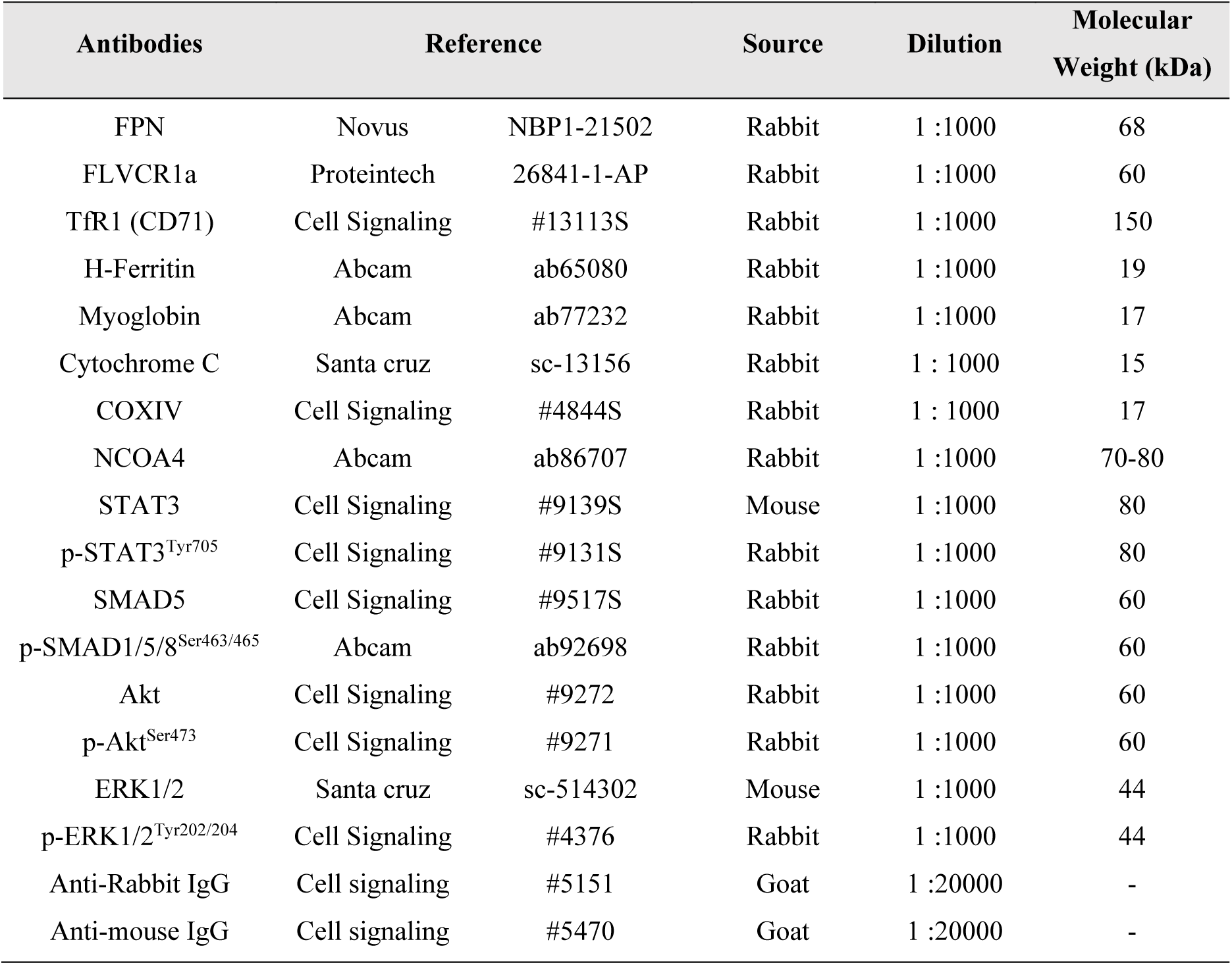
List of primary antibodies used for western blotting.

| Antibodies | Reference |  | Source | Dilution | Molecular Weight (kDa) |
| --- | --- | --- | --- | --- | --- |
| FPN | Novus | NBP1-21502 | Rabbit | 1 :1000 | 68 |
| FLVCR1a | Proteintech | 26841-1-AP | Rabbit | 1 :1000 | 60 |
| TfR1 (CD71) | Cell Signaling | #13113S | Rabbit | 1 :1000 | 150 |
| H-Ferritin | Abcam | ab65080 | Rabbit | 1 :1000 | 19 |
| Myoglobin | Abcam | ab77232 | Rabbit | 1 :1000 | 17 |
| Cytochrome C | Santa cruz | sc-13156 | Rabbit | 1 : 1000 | 15 |
| COXIV | Cell Signaling | #4844S | Rabbit | 1 : 1000 | 17 |
| NCOA4 | Abcam | ab86707 | Rabbit | 1 :1000 | 70-80 |
| STAT3 | Cell Signaling | #9139S | Mouse | 1 :1000 | 80 |
| p-STAT3<sup>Tyr705</sup> | Cell Signaling | #9131S | Rabbit | 1 :1000 | 80 |
| SMAD5 | Cell Signaling | #9517S | Rabbit | 1 :1000 | 60 |
| p-SMAD1/5/8<sup>Ser463/465</sup> | Abcam | ab92698 | Rabbit | 1 :1000 | 60 |
| Akt | Cell Signaling | #9272 | Rabbit | 1 :1000 | 60 |
| p-Akt<sup>Ser473</sup> | Cell Signaling | #9271 | Rabbit | 1 :1000 | 60 |
| ERK1/2 | Santa cruz | sc-514302 | Mouse | 1 :1000 | 44 |
| p-ERK1/2<sup>Tyr202/204</sup> | Cell Signaling | #4376 | Rabbit | 1 :1000 | 44 |
| Anti-Rabbit IgG | Cell signaling | #5151 | Goat | 1 :20000 | - |
| Anti-mouse IgG | Cell signaling | #5470 | Goat | 1 :20000 | - |

### RNA extraction and RT-qPCR

RNA was extracted from liver samples using Trizol® reagent (Sigma Aldrich), and from quadriceps samples using NucleoSpin RNA Plus kit (Macherey-Nagel) according to the manufacturers’ instructions. RNA concentrations were determined by NanoDrop spectrophotometer and reverse transcription reaction (RT) was performed (IScript RT Supermix, Bio-Rad). Then, Real-time PCR was conducted on 10 μl reaction volumes containing 4.8 μl SybrGreen SuperMix (Bio-Rad), 0.2 μl of forward and reverse primers (100 nM final) and 5 μl of cDNA (diluted at 1/40). Primer sequences are reported in Table 2. Data were analyzed using the ΔΔCT method of analysis and reference genes (*Rpl4*, *Rp119* and *Ppia*) were used for normalization.

**Table 2.**
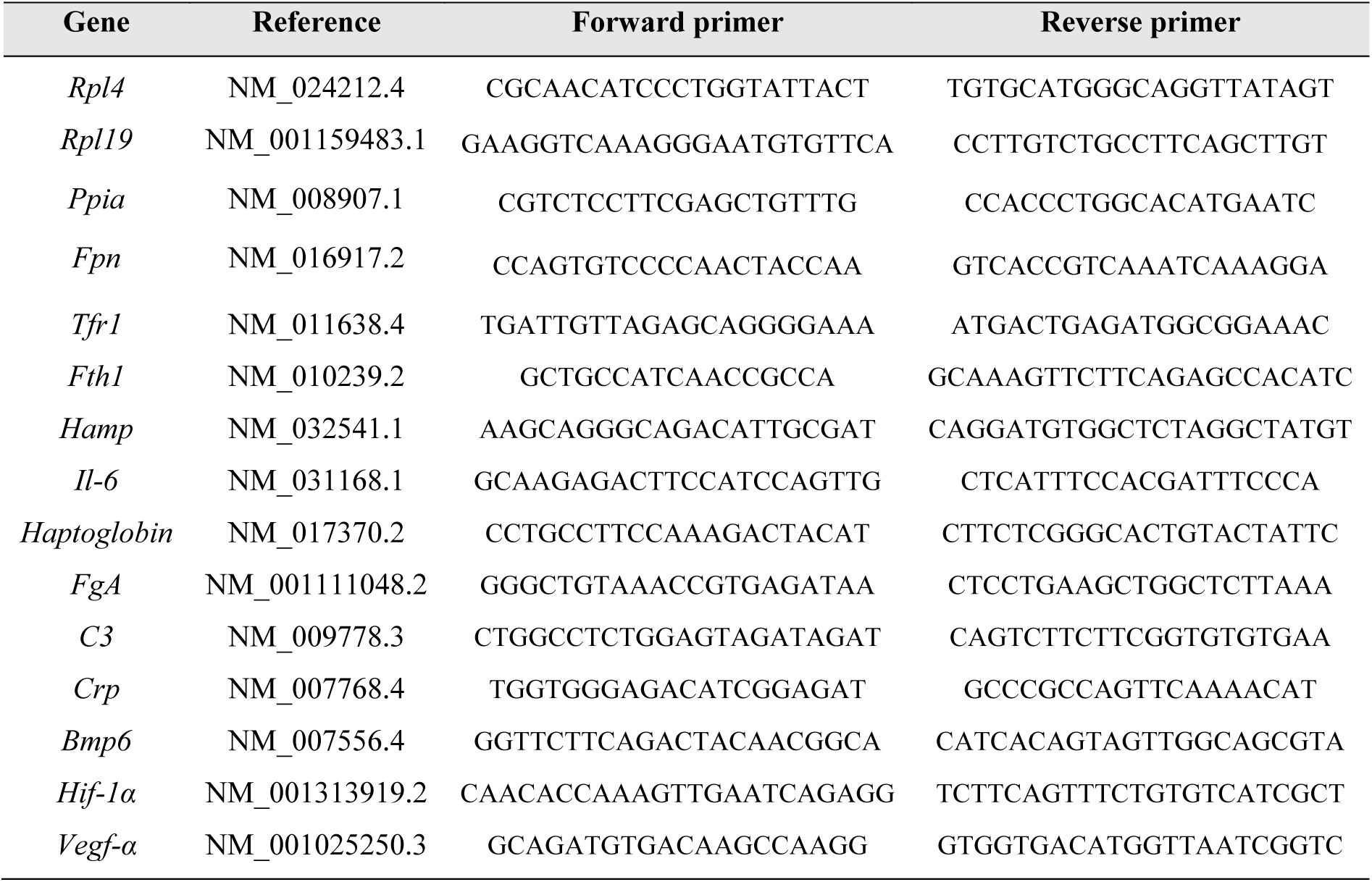
Primer design for qPCR.

### Histology analyses

Liver, spleen and tumor tissues were fixed in 4% PFA for 96h and embedded in paraffin. Serial transverse sections of 4 μm were cut from the widest part of each sample using a Microm microtome and mounted on glass slides. To visualize iron deposits, deparaffinized liver, spleen and tumor sections were stained with Perls’ Prussian Blue.

### Statistical analyses

Results are presented as individual values and mean. The normality of each distribution and homogeneity of variance were assessed with the Shapiro-Wilk and Fisher test, respectively. Groups were compared using a Student’s t-test to explore the effects of cancer cachexia. When the assumptions of normality and/or equal variance were not met, the Mann-Whitney test was used instead. To determine the effects of cancer cachexia evolution over the experimentation, two-way ANOVA were performed and followed by a Tukey’s post-hoc multiple comparison test. Spearman correlation test were used to test correlations between iron and cachexia parameters. For all analyses, statistical significance was considered when p-value < 0.05. Data were analyzed using GraphPad Prism (Version 8.0.1).

## RESULTS

### Pancreatic cancer model UN-KC-6141 induces cachexia, associated with decreased plasma iron availability

Since pancreatic cancer is the cancer most frequently associated with cachexia worldwide^4^, we first analyzed the regulation of iron metabolism in pancreatic tumor-bearing UNKC mice, compared with age-matched healthy control (CTL) mice injected with PBS.

Throughout the experiment, body weight and food intake were monitored daily to assess the onset of clinical signs of cachexia-related weight loss and anorexia (Fig. S1). Food intake was similar between UNKC and CTL mice until sacrifice, indicating the absence of anorexia in this model (Fig. S1B). Metabolic flexibility appeared to be impacted in UNKC mice over the course of the protocol, as reflected by an improvement in glucose tolerance between 3 days before cancer cell injection and 26 days after (p<0.001, Fig. S1C-E). On the day of sacrifice (i.e., 35 days post-injection), UNKC mice weighed 4.7% less than CTL mice (p=0.056, Fig.1A). Gastrocnemius and soleus muscle mass were reduced by 7.5% and 6.8% (p=0.003 and p=0.050, respectively), and this was accompanied by a 46.4% reduction of visceral fat mass in UNKC mice compared with CTL mice (p<0.001, Fig. 1B). However, quadriceps muscle mass did not significantly differ between the two groups (Fig. 1B). Together, this trend toward body weight loss, and the significant skeletal muscle and adipose tissue loss confirm that the model induced cachexia phenotype. Classification of cachexia severity revealed heterogeneous disease progression within the UNKC group, with 6 mice exhibiting early-stage cachexia, 4 moderate cachexia, and 3 severe cachexia. As inflammation is another hallmark of cancer cachexia, serum IL-6 was measured and found elevated in UNKC mice, remaining undetectable in CTL mice (p=0.001, Fig. 1F). Regarding systemic iron metabolism, circulating hepcidin increased 5-fold in UNKC mice (p<0.001, Fig. 1E), and this elevation was strongly correlated with the rise in serum IL-6 concentrations (r=0.70, p<0.001). In addition, transferrin saturation decreased compared to CTL mice (-9.6%, p<0.001, Fig. 1D), supporting reduced plasma iron availability, with no concomitant change in hematocrit (Fig. 1C).

**Figure 1.**
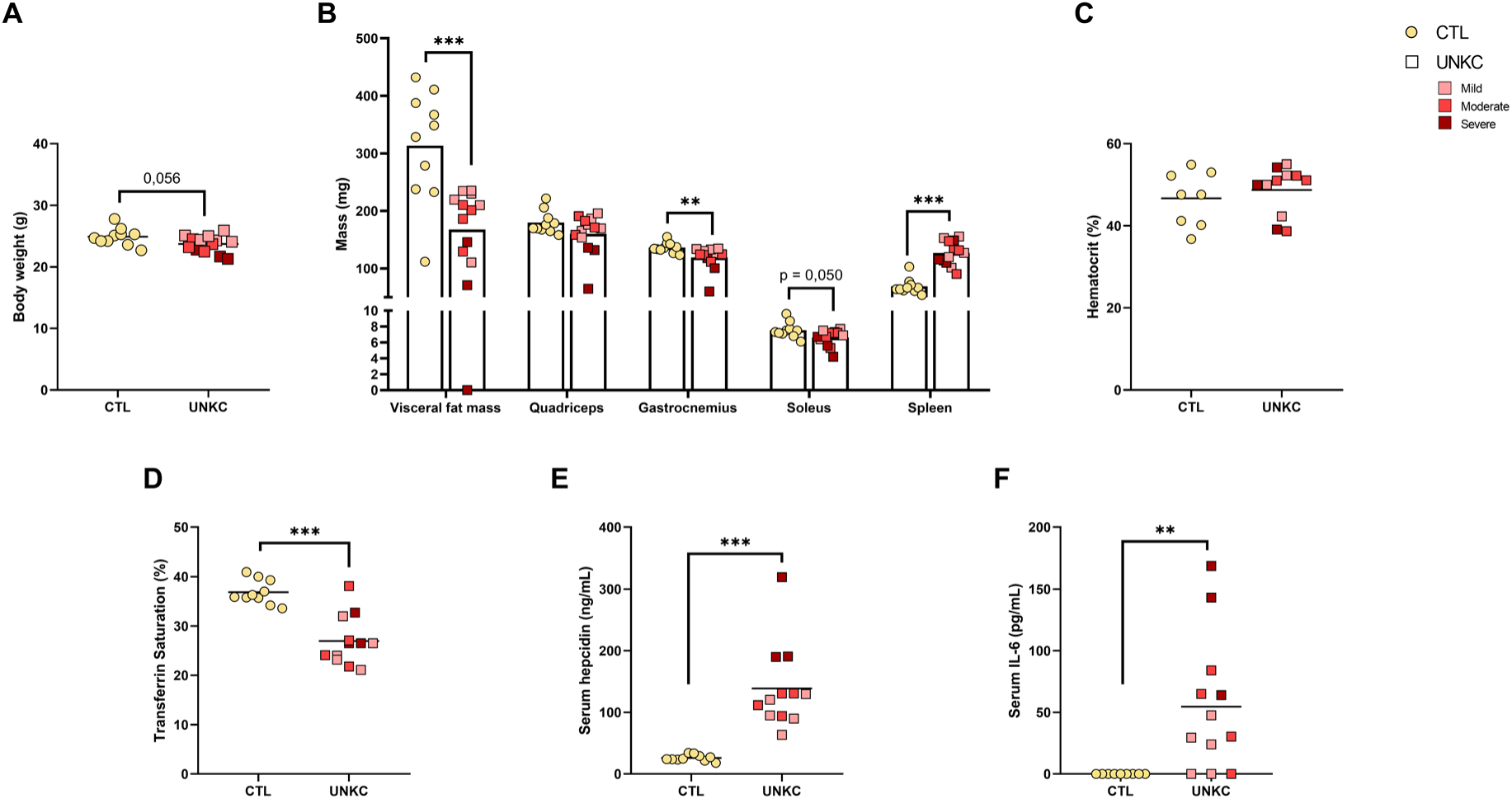
UNKC pancreatic cancer is associated with cachexia, reduced iron availability and increased serum hepcidin levels. (A) Mice body weight without tumor (g) (B) Visceral fat, quadriceps, gastrocnemius, soleus and spleen mass (C) Hematocrit (D) Transferrin saturation (E) Serum hepcidin levels (F) Serum IL-6 levels. Serum IL-6 values below the lowest point of the standard curve (7.8 pg/mL) were considered undetectable and were assigned a value of 7.8 pg/mL for statistical analysis using the Mann-Whitney test. Data were represented as individual value and mean. Significance was checked using Mann-Whitney tests: **p < 0.01, ***p < 0.001 (CTL, n= 10; UNKC, n=13).

### Systemic inflammation associated with cachectic pancreatic cancer upregulates hepatic hepcidin synthesis, promoting splenic iron sequestration

Under cancer cachexia conditions, the liver is one of the main organs affected, displaying metabolic reprogramming that alters hepatic function. It also plays a central role in iron metabolism, ensuring body iron storage and acting as the main site of hepcidin synthesis^15,16^. The liver is therefore rapidly affected in cases of iron excess or deficiency. For these reasons, we explored the expression of iron metabolism-related genes and proteins in the liver of UNKC mice.

Compared with CTL mice, liver iron concentrations (LIC), as well as FtH, FPN and TfR1 proteins remained unchanged in the UNKC group (Fig. 2A-D). However, the 2-fold increase in *Hamp* mRNA encoding hepcidin (p<0.001, Fig. 2E), which strongly correlated with circulating hepcidin concentrations (r=0.81, p<0.001), reflects an upregulation of hepatic hepcidin synthesis in pancreatic tumor-bearing mice. We then analyzed the expression of hepatic genes and proteins involved in the iron- and inflammation-related pathways regulating hepcidin synthesis^9^ (Fig. 2F-H). In UNKC mice, we observed significant increases in the mRNA expression of all pro-inflammatory markers studied (i.e., *Haptoglobin*, *Fibrinogen A*, *C3*, *Il-6*, *Crp*, encoding acute-phase proteins, Fig. 2H), together with a 2-fold increase in phosphorylated STAT3^Tyr705^ protein levels (p<0.001, Fig. 2F and 2G), the key transcription factor driving *Hamp* expression under pro-inflammatory conditions^31^. Conversely, *Bmp6* mRNA, which is overexpressed upon activation of the iron-dependent pathway of hepcidin synthesis^32^, was expressed at lower levels in UNKC than CTL mice (-42%, p<0.001, Fig. 2H), although the phosphorylated SMAD1/5/8^Ser463/465^ protein associated with this pathway showed a slight trend toward increased expression (p=0.077, Fig. 2F and 2G).

**Figure 2.**
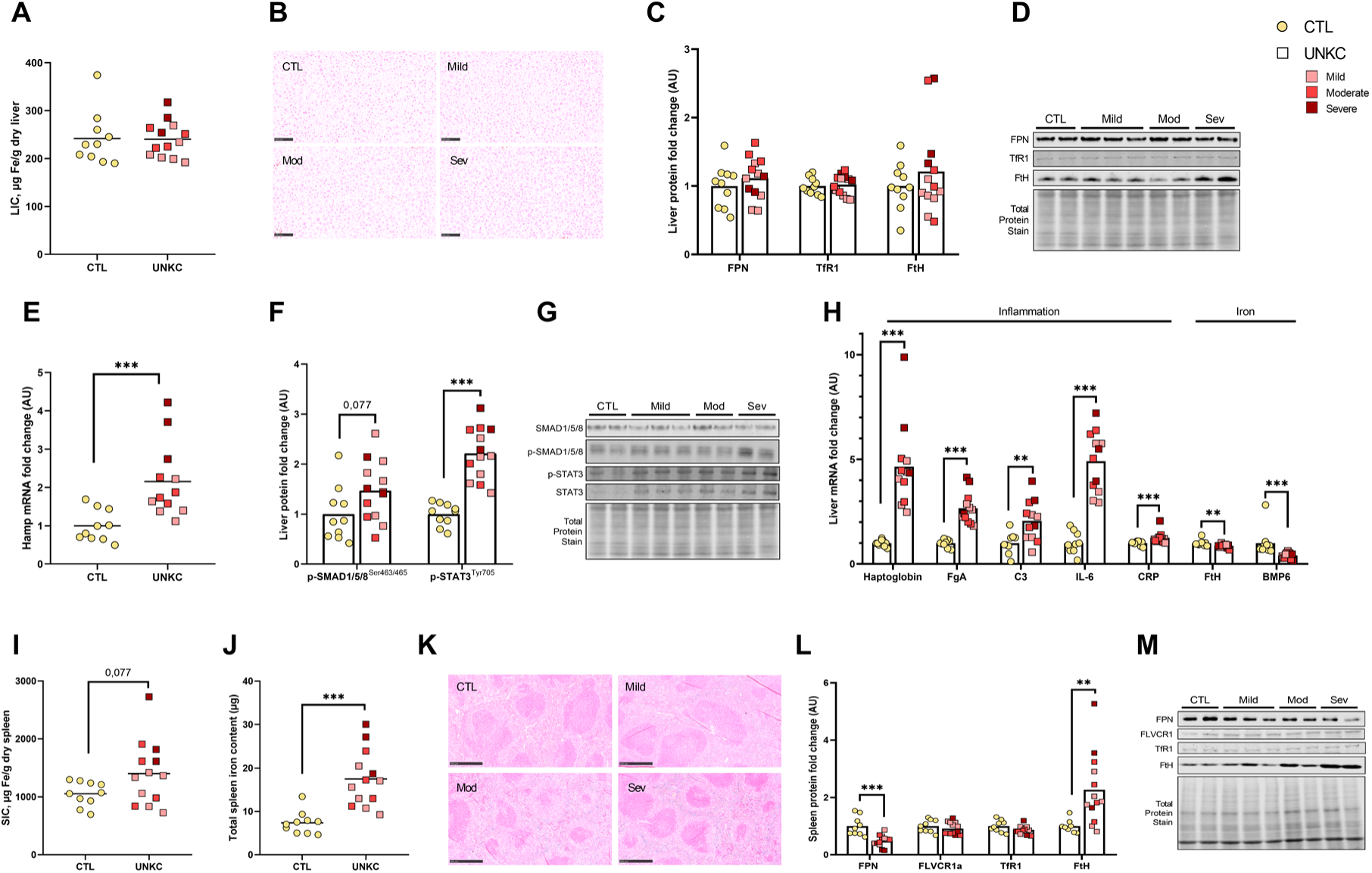
Hepcidin synthesis is upregulated by inflammation and associated to spleen iron sequestration in UNKC mice. (A) Liver iron concentrations (B) Reference image of liver histological blue Perls coloration (C) Expression levels of FPN, TfR1 and FtH proteins and (D) its western blot reference images (E) Expression level of *Hamp* mRNA in liver (F) Expression levels of p-SMAD1/5/8^Ser463/465^ and p-STAT3^Tyr705^ proteins and (G) its western blot reference images (H) Expression levels of pro-inflammatory and iron related genes mRNA (I) Spleen iron concentrations (J) Spleen total iron pool (K) Reference image of spleen histological blue Perls coloration (L) Expression levels of FPN, FLVCR1a, TfR1 and FtH proteins, (M) Spleen Western blot reference images. Data were represented as individual value and mean. Significance was checked using Mann-Whitney tests: **p < 0.01, ***p < 0.001 (CTL, n= 10; UNKC, n=13).

Like the liver, the spleen is a major organ in iron metabolism, notably involved in red blood cell recycling and capable of storing large amounts of iron when present in excess^8^. Moreover, given its high density of macrophages, the cell type expressing the highest levels of FPN, the spleen is particularly responsive to hepcidin action. Here, we observed that pancreatic cancer tends to increase spleen iron concentrations (SIC) in UNKC mice (p=0.077, Fig. 2I and 2K), despite a marked increase in spleen weight (+95.5%, p<0.001, Fig. 1B). Consequently, total splenic iron content was approximately doubled compared to CTL mice (+134%, p<0.001, Fig. 2J and 2K). This was accompanied by increased splenic FtH protein expression (+90.7%, p=0.001), whereas FPN protein levels decreased (-42.4%, p<0.001), likely in response to hepcidin, together indicating splenic iron sequestration (Fig. 2L and 2M). Consistent with this finding, Perl’s staining showed iron to be more abundant in the red pulp, where macrophages are predominantly located (Fig. 2K).

### Skeletal muscle adapts to preserve iron homeostasis under pancreatic cancer cachexia

Given the central role of skeletal muscle in cachexia and iron metabolism, we then explored in the UNKC model the regulation of the main iron-related genes and proteins in two muscles, one atrophied (i.e., gastrocnemius) and one non-atrophied (i.e., quadriceps). In gastrocnemius muscle, iron concentrations remained unchanged (Fig. 3A), but total iron content significantly decreased (-12.9%, p=0.017, Fig. 3B), due to the concomitant muscle atrophy (Fig. 1B). Regarding iron transport proteins, we observed no significant modulation in the expression of the iron and heme export proteins FPN and FLVCR1a, and only a slight trend toward increased levels of the iron import protein TfR1 (p=0.091, Fig. 3C and 3D). However, FtH protein expression significantly increased (+86.6%, p=0.010, Fig. 3C and 3D). We also investigated the protein expression of key hemoproteins including myoglobin, cytochrome c (CYSC), and cytochrome c oxidase (COXIV), which incorporate iron through heme and are essential to numerous biological functions such as oxygen binding and oxidative phosphorylation^33,34^. Here, we observed that myoglobin and COXIV expression were significantly increased in the atrophied gastrocnemius muscle (+38%, p=0.044 and +33%, p=0.002, respectively), with a similar trend for CYSC (+17%, p=0.069, Fig. 3C and 3D). In quadriceps muscle, iron concentrations and total iron content were similar to CTL mice (Fig. 3E and 3F). Proteins and genes involved in iron transport, storage and utilization remained unchanged, with the exception of myoglobin, whose expression increase in UNKC mice (+34%, p=0.031, Fig. 3G-I).

**Figure 3.**
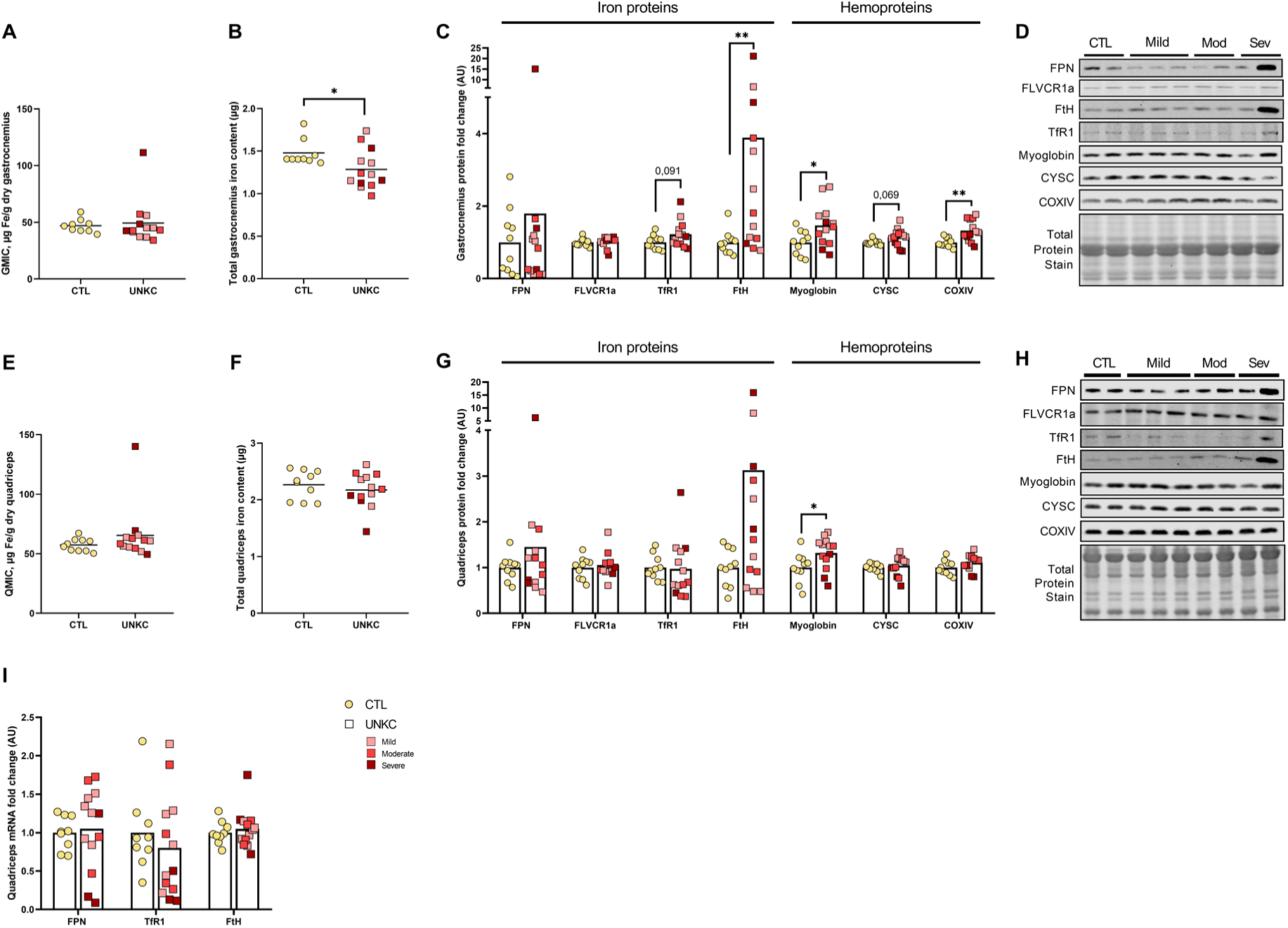
Skeletal muscles maintain iron concentrations despite muscle atrophy in UNKC mice. (A) Gastrocnemius muscle iron concentrations (B) Gastrocnemius total iron pool (C) Expression levels of iron metabolism related proteins in gastrocnemius (D) Gastrocnemius Western blot reference images (E) Quadriceps muscle iron concentrations (F) Quadriceps total iron pool (G) Expression levels of iron metabolism related proteins in quadriceps (H) Quadriceps Western blot reference images (I) Expression levels of *FPN*, *TfR1* and *FtH* mRNA in quadriceps. Data were represented as individual value and mean. Significance was checked using Mann-Whitney tests: *p < 0.05, **p < 0.01 (CTL, n= 10; UNKC, n=13).

### Tumor iron metabolism in UNKC pancreatic cancer

Iron concentrations, as well as the expression of iron metabolism regulatory proteins (i.e., FPN, TfR1, H-ferritin, NCOA4) and proliferation-related proteins (p-Akt, p-ERK), were determined in tumors from UNKC mice. Perls’ staining was also performed for qualitative analysis (Supplemental Fig. S2). Here, we reported that tumor iron concentration was negatively associated with tumor mass (r=-0.56, p=0.05, Supplemental Fig. S2D), although larger tumors contained higher total iron content (r=0.824, p<0.001). Expression of the iron import protein TfR1 in the tumor also showed a trend toward association with tumor mass (r=0.539, p=0.06), and is notably positively correlated with the phosphorylation levels of ERK and Akt (r=0.578, p=0.042 and r=0.614, p=0.028, respectively), as well as with body weight loss (r=-0.732, p=0.006) and gastrocnemius muscle mass loss (r=-0.669, p=0.015, Supplemental Fig. S2G and S2H).

### The colorectal cancer model Apc^Min/+^ induces cachexia and anemia, associated with organ iron depletion despite reduced hepcidin synthesis

Colorectal cancer, like pancreatic cancer, is frequently associated with cachexia, and is particularly characterized by the presence of intestinal polyps prone to bleeding. Consequently, anemia prevalence is elevated in this type of cancer^6,7^. We therefore explored the regulation of iron metabolism in the *Apc*^Min/+^ murine model of colorectal cancer^27^, analyzing the liver, spleen, and quadriceps muscle of these mice (Fig. 4 and 5).

**Figure 4.**
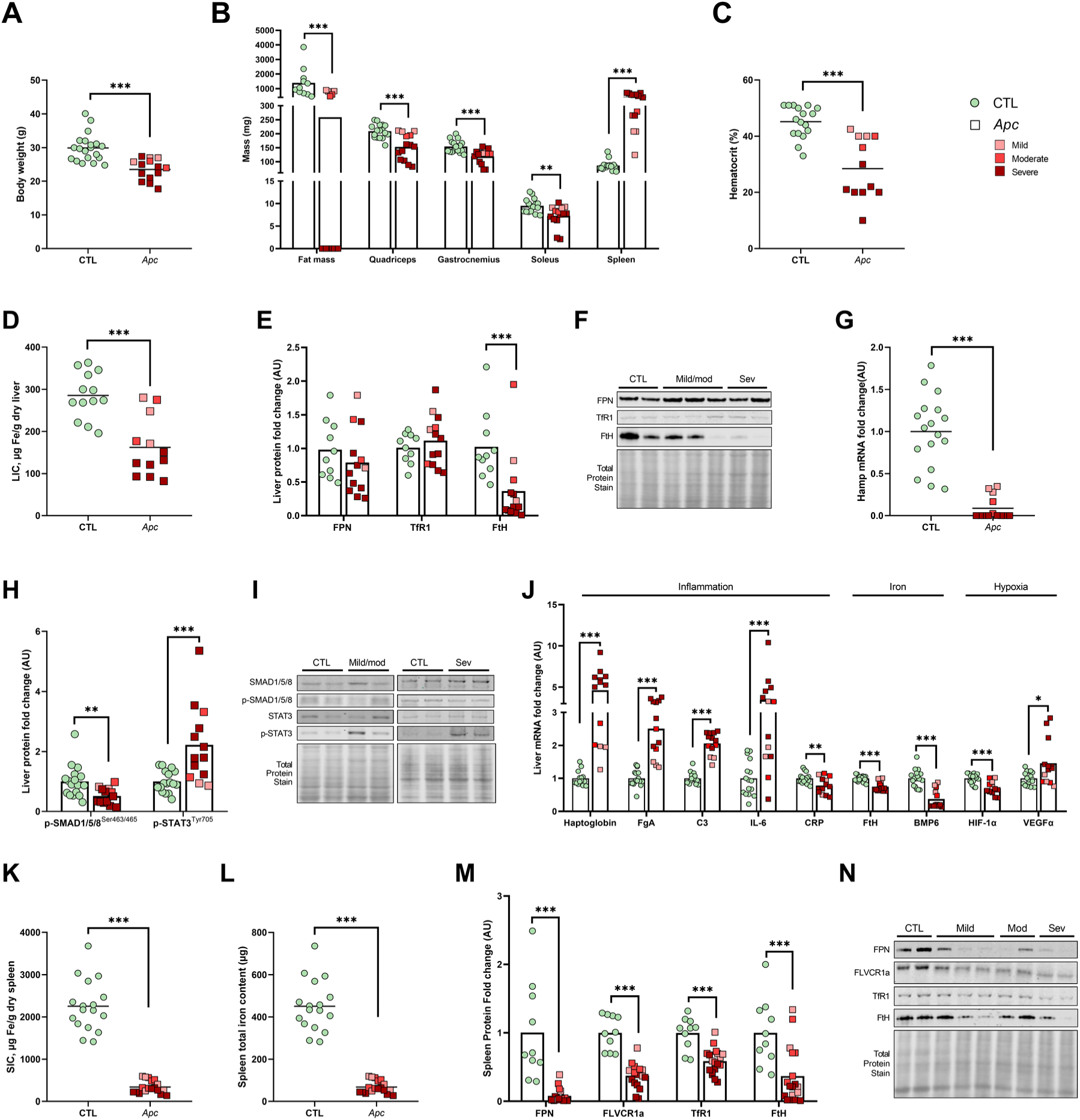
*Apc*^Min/+^ colorectal cancer is associated with cachexia, anemia, reduced iron levels in liver and spleen, and downregulated hepcidin synthesis. (A) Mice body weight (g) (B) Fat, quadriceps, gastrocnemius, soleus and spleen mass (C) Hematocrit (D) Liver iron concentrations (E) Expression levels of FPN, TfR1 and FtH proteins and (F) its western blot reference images (G) Expression level of *Hamp* mRNA in liver (H) Expression levels of p-SMAD1/5/8^Ser463/465^ and p-STAT3^Tyr705^ proteins and (I) its western blot reference images (J) Expression levels of pro-inflammatory, iron and hypoxia related genes mRNA (K) Spleen iron concentrations (L) Spleen total iron pool (M) Expression levels of FPN, FLVCR1a, TfR1 and FtH proteins (N) Spleen Western blot reference images. Data were represented as individual value and mean. Significance was checked using Mann-Whitney tests: *p < 0.05, **p < 0.01, ***p < 0.001 (CTL, n= 24; *Apc*, n=24).

**Figure 5.**
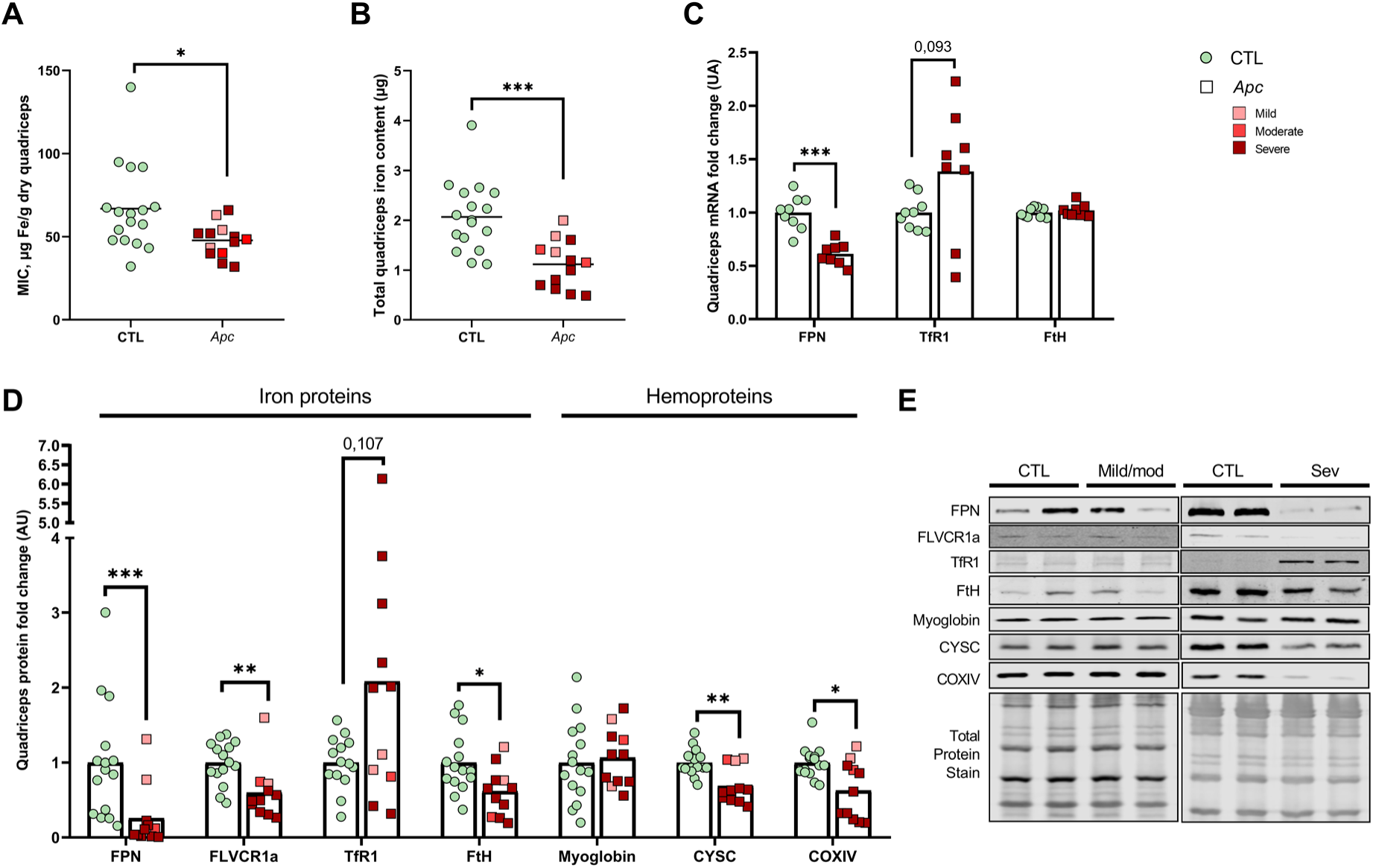
Skeletal muscle modulates its iron distribution through cellular regulation in *Apc* mice. (A) Quadriceps muscle iron concentrations (B) Quadriceps total iron pool (C) Expression levels of *FPN*, *TfR1* and *FtH* mRNA in quadriceps (D) Expression levels of iron metabolism related proteins in quadriceps (E) Quadriceps Western blot reference images. Data were represented as individual value and mean. Significance was checked using Mann-Whitney tests: *p < 0.05, **p < 0.01, ***p < 0.001 (CTL, n= 17; *Apc*, n=13).

As in the UNKC model, body weight loss (-21.4%, p<0.001), together with significant fat mass and muscle mass loss (i.e., quadriceps, gastrocnemius, soleus), confirm cachexia induction in the *Apc* model (Fig. 4A and 4B). But on contrary to UNKC model, hematocrit was markedly decreased in *Apc* mice compared with their healthy littermates (-43.5%, p<0.001), supporting the presence of anemia (Fig. 4C). In the liver, iron concentrations decreased (-48.4%, p<0.001, Fig. 4D) and were associated with reduced FtH protein levels (-85.2%, p<0.001, Fig. 4E and 4F). In contrast to UNKC mice, we observed a drop in hepatic *Hamp* mRNA expression in *Apc* mice (-99.9%, p<0.001, Figure 4G), despite the pro-inflammatory state of the animals, reflected by increased hepatic mRNA levels of inflammatory markers (*Haptoglobin*, *FibrinogenA*, *C3* and *Il-6*, p<0.001, Fig. 4J) and enhanced phosphorylated STAT3^Tyr705^ protein expression (p<0.001, Fig. 4H and 4I). As the *Apc* model is associated with anemia, we explored several hepatic hypoxia-related genes, including *Hif-1α* and *Vegf-α*, a HIF-1α-responsive gene that promotes angiogenesis^35^. Hepatic *Vegf-α* mRNA expression increased in *Apc* mice (+36.4%, p=0.024, Fig. 4J). Regarding the genes and proteins involved in the iron-dependent pathway of hepcidin synthesis, *Bmp6* mRNA expression significantly decreased (-79.7%, p<0.001, Fig. 4J), as well as hepatic levels of phosphorylated-SMAD1/5/8^Ser463/465^ protein (-44%, p=0.003, Fig. 4H and 4I). Despite the marked decrease in hepatic hepcidin synthesis, splenic FPN levels were significantly reduced (-94.2%, p<0.001), and the spleen is markedly depleted of iron (-84.9%, p<0.001, Figures 4K, 4L). The heme exporter FLVCR1a, the iron import protein TfR1, and the storage FtH protein were also strongly under-expressed in the spleen of *Apc* mice (p<0.001, Figures 4M and 4N).

### Skeletal muscle iron regulation diverges from systemic signals to protect itself against iron deficiency

In the atrophied quadriceps muscle (Fig. 4B) of *Apc* mice, iron concentrations also decreased compared with muscle from CTL mice (-24.1%, p=0.012), contributing to a significant reduction in total muscle iron content (-42.3%, p<0.001, Fig. 5A and 5B). Concomitantly, our analyses revealed a decrease in both FPN mRNA and protein levels (-40.4%, p<0.001 and -87.9%, p<0.001, respectively), associated with a trend toward increased TfR1 mRNA and protein levels, suggesting the involvement of cellular regulation via the IRE/IRP pathway in the atrophied quadriceps muscle (Fig. 5C-E). Moreover, the expression of the heme export protein FLVCR1a, of FtH, and of the hemoproteins CYSC and COXIV was reduced, supporting downregulation of muscle iron export, storage, and utilization in response to cellular iron deficiency. Only myoglobin protein levels appeared to remain stable in *Apc* mice (Fig. 5D and 5E).

## DISCUSSION

In the present study, we investigated the regulation of systemic and cellular iron metabolism in two distinct murine models of cancer-induced cachexia, either associated with or without anemia. Although both models induced cachexia, we report distinct patterns of systemic iron regulation. In the UNKC pancreatic cancer model (without anemia), the pro-inflammatory state promotes hepatic hepcidin synthesis and iron redistribution towards splenic sequestration. In the *Apc*^Min/+^ colorectal cancer model, associated with anemia, mice exhibit systemic iron depletion across the liver, spleen and skeletal muscle, and the associated iron deficiency and/or hypoxia signals appear to override inflammatory signals, driving hepcidin downregulation. In both models, however, iron handling in atrophied skeletal muscle appears to escape systemic hepcidin control, being instead primarily driven by local cellular iron status.

### Murine models of cachectic pancreatic and colorectal cancers both induce iron redistribution, but through divergent regulatory mechanisms, including opposite hepcidin responses

In UNKC mice, reduced plasma iron availability occurs alongside inflammation-driven hepcidin upregulation, consistent with the well-established capacity of the IL-6/STAT3 inflammatory pathway to induce hepatic hepcidin synthesis^9,17,31^. Hepatic iron stores remain unaffected, whereas the spleen accumulates iron within its macrophage-rich red pulp, as a result of reduced FPN expression. Such pattern is consistent with hepcidin-driven macrophage iron sequestration previously described in inflammatory anemia^17^ or muscle disuse^22,24,25^. Inflammation therefore appears to be the principal driver of hepcidin regulation, and consequently of iron redistribution in this pancreatic cancer model.

Conversely, in the *Apc* model associated with anemia – evidenced by decreased hematocrit and supported by the literature^26^ – our results show hepcidin downregulation despite the inflammatory context. This coincides with markers of hepatic hypoxia, together with reduced activity of the iron-sensing BMP6/SMAD pathway and marked hepatic and splenic iron depletion. These results indicate that, in this context combining cachexia and anemia, hypoxia and/or iron-deficiency signals override the inflammatory drive on hepatic hepcidin synthesis, a hierarchy consistent with that previously reported among these signals in other pathophysiological contexts^36–38^. Despite this hepcidin downregulation, FPN levels are substantially decreased in the spleen of *Apc* mice. FLVCR1a, TfR1, and FtH protein levels are also decreased, limiting iron export, import, and storage in the iron-depleted spleen, and likely reflecting a functional collapse of the spleen’s capacity to handle iron under these conditions of severe iron deficiency. Altogether, these results show that systemic iron redistribution during cachexia is cancer type-specific, and appears substantially influenced by the presence or absence of concomitant anemia. Notably, both models share a decrease in iron availability, consistent with the high prevalence of iron deficiency reported across cancer types^39^.

### Whatever the cachectic cancer model, atrophied skeletal muscle appears to manage its iron according to local needs, with limited influence from systemic hepcidin action

Skeletal muscle, whose mass and function are particularly altered by cancer cachexia, is among the organs with the highest iron demand, owing to its energy requirements and the presence of myoglobin within muscle fibers^33^. However, the regulation of muscle iron metabolism remains poorly described, particularly in the context of cancer cachexia, even though skeletal muscle constitutes the primary target of clinical management^1,4^. The analyses of gastrocnemius and quadriceps muscles reveal a differential impact of the two cachectic cancer types on muscle iron regulation, which appear to be more markedly affected in *Apc* mice, when cachexia is associated with anemia.

In the UNKC model, the gastrocnemius muscle maintains its iron concentration despite atrophy, by releasing part of its total iron pool. Concomitantly, increased expression of FtH and the hemoproteins myoglobin, CYSC and COXIV could reflect a cellular strategy to limit the expansion of the labile iron pool (LIP) by channeling iron into non-toxic, functional forms (i.e. heme), potentially also compensating for the mitochondrial dysfunction associated with cachexia^5^. By contrast, muscle iron metabolism in the non-atrophied quadriceps of UNKC mice is not altered by the systemic environment of pancreatic cancer cachexia. It nonetheless shows increased myoglobin levels, suggesting a regulatory trend similar to that observed in the atrophied gastrocnemius muscle. In both atrophied and non-atrophied muscles, FPN levels remain unchanged despite elevated circulating hepcidin. This may reflect a concomitant compensatory cellular regulation of FPN synthesis in muscle, counteracting its hepcidin-mediated degradation. The IRE/IRP system could be involved in this compensatory regulation, as supported by increased FtH protein levels in the gastrocnemius; however, the absence of decreased TfR1 mRNA (in the quadriceps) and protein levels (in both muscles) does not allow this involvement to be confirmed. Alternatively, if these maintained FPN levels are not explained by cellular regulation, muscle FPN may not respond to systemic hepcidin, or at least is not degraded by it. It cannot be excluded that it is simply inactivated rather than degraded, as suggested by more recent findings^16^. In UNKC mice, skeletal muscle thus seems to adapt in order to limit the toxic LIP through the release of part of its iron pool, the storage of iron via FtH and the synthesis of heme-containing proteins to incorporate iron.

In *Apc* mice, muscle iron distribution in the quadriceps is significantly affected by cachexia and anemia-related iron-deficiency context. This atrophied muscle is also iron-depleted, as reflected by decreased iron concentrations and total iron content. Despite hepcidin downregulation, FPN expression decreases, reflecting the predominance of local cellular regulation. Indeed, this modulation, together with the increase in TfR1 mRNA, is consistent with an activation of the IRE/IRP system, which is engaged under iron deficiency. The membrane protein FLVCR1 is also downregulated, suggesting sequestration of iron in its ferrous and heme-bound forms. Finally, the levels of FtH protein and hemoproteins CYSC and COXIV decrease, further supporting the severity of iron deficiency in the atrophied quadriceps.

Regardless of the model, muscle iron metabolism does not appear to respond to systemic hepcidin-mediated regulation, or at least is substantially less responsive than other organs expressing different cell types. This independence of skeletal muscle from systemic hepcidin action points to cellular IRE/IRP-mediated regulation as the most likely explanation, although this remains to be confirmed. This local regulation nonetheless differs markedly between the two cancer types: skeletal muscle preserves its iron concentration in UNKC mice, redistributing iron toward storage and heme-protein incorporation, whereas it favors iron retention in *Apc*^Min/+^ mice, consistent with the severe systemic iron depletion associated with anemia in this model.

This divergence suggests that cancer type, and probably the presence of associated anemia, shapes the direction of muscle iron handling. This cancer type-specificity is further illustrated by the C26 model of cachectic colorectal cancer, in which previous studies reported yet another distinct muscle iron regulation pattern, with increased H-ferritin and protein-bound iron together with TfR1 downregulation in the atrophied gastrocnemius^40^.

### The tumor may take up part of circulating iron to fuel its own growth, potentially aggravating cachexia

Within this cachexia-associated iron redistribution, the tumor itself may play a significant role. Cancer cells have been described as displaying an “iron addiction”, enabling them to modulate their iron metabolism to increase intracellular iron levels. Tumors notably display increased TfR1 expression together with low FPN levels, expanding the labile iron pool and thereby fueling their proliferation^41,42^. Our findings in the tumors of UNKC mice are consistent with these previous observations and raise the possibility of a link between tumor size, cachexia progression (i.e., body weight and muscle mass loss), and tumor iron import (i.e., TfR1 upregulation). However, the number of animals per cachexia severity grade was too small to establish clear associations, and this hypothesis warrants further investigation.

## CONCLUSION

In summary, our data reveal that (1) there is no common regulatory signature of iron metabolism across two different cachexia conditions, (2) iron regulation is driven more by cancer type (notably, whether it is associated with anemia or not) than by cachexia severity, and (3) muscle iron metabolism is impaired in cachexia associated with anemia, but not necessarily in its absence despite atrophy. Iron deficiency appears to be a common feature across the different types of cachectic cancer, but its severity varies considerably between models, differentially affecting iron distribution across organs. Considering these divergences, the clinical management of iron metabolism in cachectic patients should be guided by the underlying cancer type.

## AUTHOR CONTRIBUTIONS

M.A., D.F., A.R. and F.D. designed research studies. M.A., A.R. and F.D. supervised the study. M.A., P.G., L.O., B.M., M-L.I., M.R., A.R. and F.D. performed experiments. Statistics analyses was provided by M.A. M.A., F.D. and A.R. analyzed data, interpreted results of experiments and wrote the paper. All authors revised the manuscript. All authors have read and agreed to the published version of the manuscript.

## ACKNOWLEDGEMENTS

The authors thank Pascale Bellaud (H2P2 facility, University Rennes 1, Rennes) for technical help with histological analyses. The authors also thank Olivier Loréal for valuable scientific discussions during the analysis of the data.

## SUPPLEMENTAL DATA

**Figure S1.**
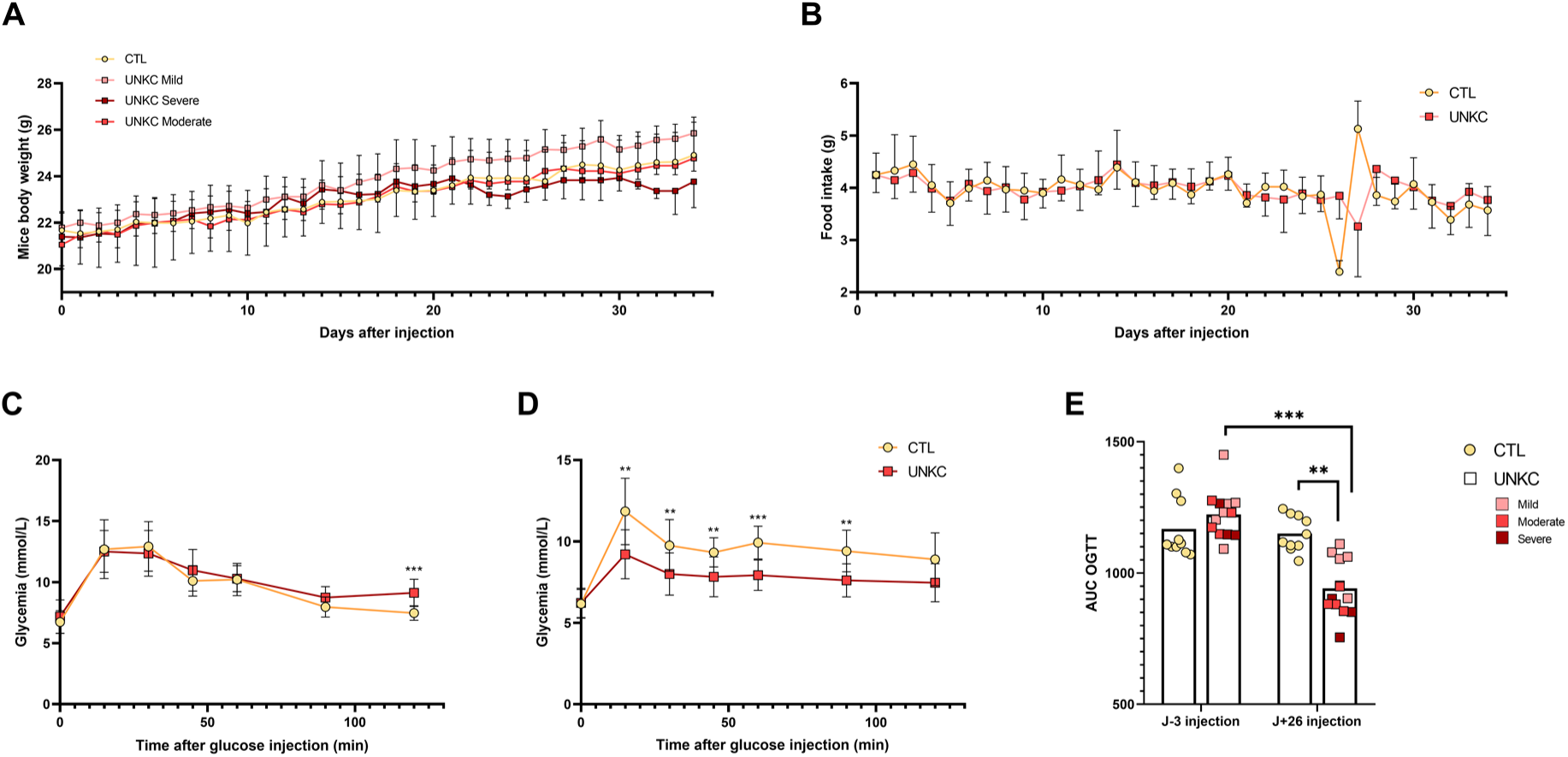
Body weight, food intake and glucose tolerance in healthy and UNKC mice during experimentation. (A) Body weight evolution including tumor (UNKC mild, n=6; UNKC moderate, n=4; UNKC severe, n=3). (B) Food intake evolution (C) Oral glucose tolerance test (OGTT) 3 days before cancer cells injection (D) Oral glucose tolerance test (OGTT) 26 days after cancer cells injection (E) Areas under the curve before and after tumor cell injection. Data are represented as mean ± SD or with individual values. Significance was checked using two-way ANOVA test and Tukey’s post-hoc test: *p < 0.05, **p < 0.01, ***p < 0.001 (CTL, n=10; UNKC, n=13).

**Figure S2.**
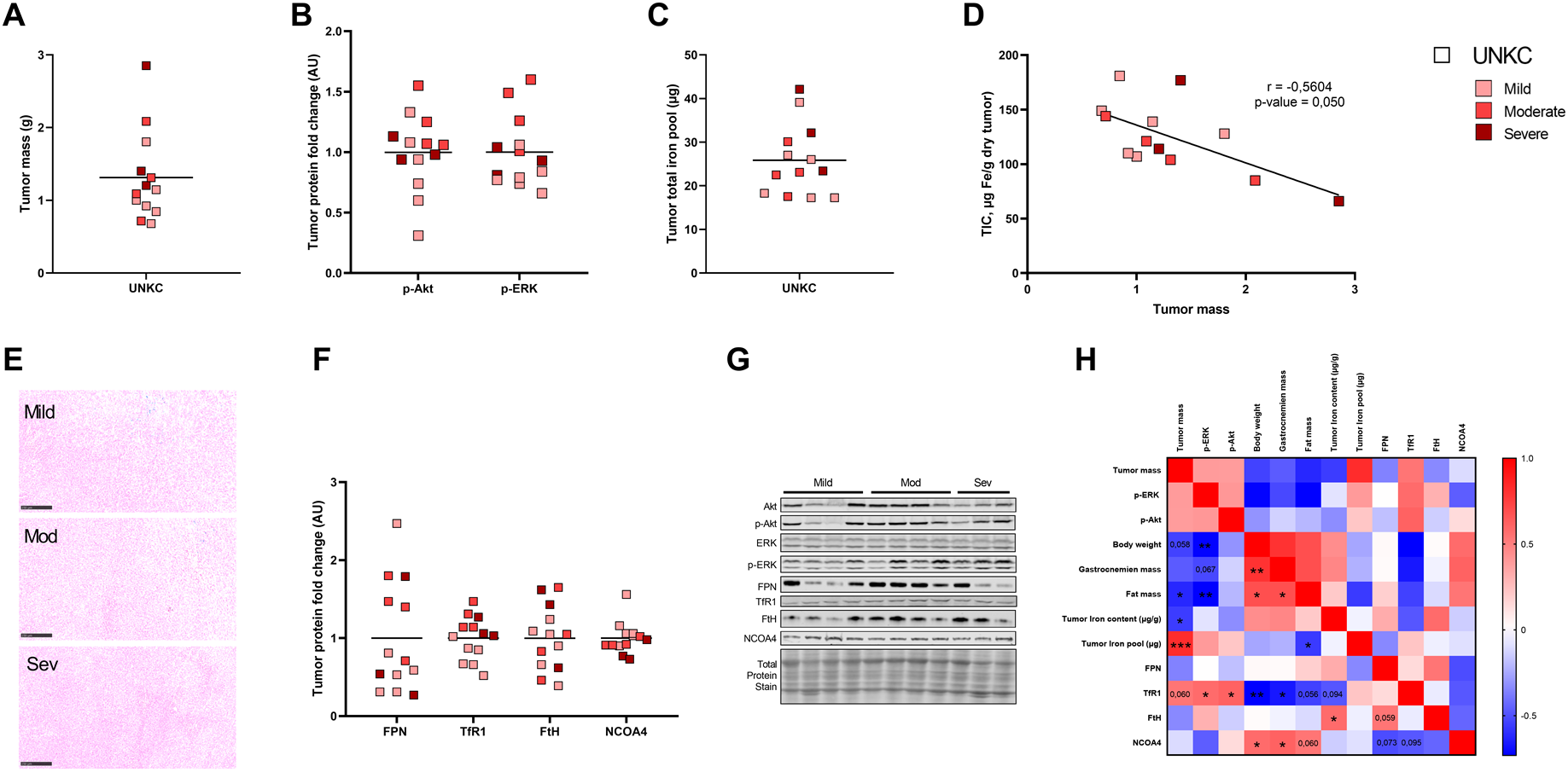
Tumor iron metabolism in UNKC mice. (A) Tumor mass (B) Expression levels of p-Akt and p-ERK proteins (C) Tumor total iron pool (D) Correlation between tumor mass and tumor iron concentrations (E) Reference image of tumor histological blue Perls coloration (F) Expression levels of iron metabolism related proteins in tumor (G) Tumor western blot representative images (H) Heatmap representing correlations between tumor growth and tumor iron metabolism parameters. Data are represented as individual values and mean and were analyzed with Spearman correlation test: *p < 0.05, **p < 0.01, ***p < 0.001.

